# PigGPA: A Domain-Constrained Agent for Porcine Genomic Breeding with Verifiable Tool Execution

**DOI:** 10.64898/2026.10.18.759908

**Authors:** Chongxiao Liang, Yanlin Zhang, Wenkang Wei, Zishuai Wang, Kui Li

## Abstract

Large language model agents can plan and execute scientific analyses, but successful task completion does not establish whether analytical decisions remain consistent across independent sessions or whether computations follow the parameters specified beforehand. We investigated these limitations using four genomic analysis tasks. General-purpose agents showed limited consistency in analytical decisions across repeated sessions (57.5–58.8%). Incorporating domain knowledge improved analytical performance, with reusable skills increasing mean evaluation scores from 73.44 to 89.06/100 , but did not resolve decision variability or ensure agreement between specified and executed parameters. To address these limitations, we developed an approach combining domain-constrained analytical skills with call-level execution contracts. Each tool invocation is linked to a pre-execution parameter declaration through SHA-256 hashing, with actual command-line arguments and execution records subsequently checked against the declared parameters. We implemented this approach in PigGPA, an autonomous agent for porcine genomic breeding that integrates statistical genetics, sequence-based functional prediction and multi-omics annotation. PigGPA completed all benchmark tasks with full agreement between declared and executed parameters, while its core quantitative-genetic results closely matched established software (r = 0.999–1.000). An application to loin muscle depth further demonstrated an integrated workflow from genetic association analysis to functional interpretation. These findings show that domain knowledge improves analytical performance, while direct verification of tool execution provides an additional basis for reproducible scientific computation.

## 1. Introduction

Large language model (LLM) agents are extending artificial intelligence from generating scientific content and code to planning analyses, executing computational tools and adapting subsequent actions based on intermediate results. The integration of reasoning and tool use, exemplified by the ReAct framework (Yao et al., 2023), has enabled applications ranging from autonomous chemical experimentation to collaborative scientific research (Boiko et al., 2023; Swanson et al., 2025). As these systems assume greater responsibility for scientific computation, their reliability cannot be established solely by successful task completion or the apparent correctness of final outputs. Two distinct questions arise: whether agents make consistent analytical decisions across independent sessions, and whether the computations they execute follow the methods and parameters specified beforehand. An agent may repeatedly propose a valid analysis yet omit essential steps during execution, modify parameters without updating its declarations, or report successful completion despite an unsuccessful tool invocation. Distinguishing the consistency of analytical decisions from agreement between declared and executed procedures is therefore essential for evaluating autonomous scientific analysis.

Genomic breeding provides a demanding setting for investigating these problems because it combines established statistical methods, heterogeneous datasets and interdependent computational procedures. Since the introduction of genomic selection (Meuwissen et al., 2001), genomic prediction has become integral to livestock genetic evaluation, including commercial pig breeding (Knol et al., 2016). Advances in reference genomes and functional genomics have further expanded these analyses from genetic association and prediction to the interpretation of candidate regulatory variants (Groenen et al., 2012; Pan et al., 2021; Teng et al., 2024). Such workflows require coordinated decisions regarding genotype quality control, population structure, statistical models, relationship matrices and validation strategies, often followed by sequence-based functional prediction and multi-omics annotation. Errors in these decisions or their implementation can affect downstream inference without necessarily preventing the generation of plausible results. Recent approaches have sought to improve analytical reliability by incorporating specialized knowledge and reusable computational tools. BioAgents and Paper2Agent demonstrated approaches to knowledge-assisted bioinformatics analysis and the reuse of published computational methods (Mehandru et al., 2025; Miao et al., 2026), while our previously developed BOLE framework incorporated domain knowledge and analytical scripts into a multi-agent system for genomic breeding (Wang et al., 2026). Although these approaches can improve method selection and task completion, access to appropriate knowledge and tools does not establish whether individual analyses were executed according to their declared procedures.

Research on reproducible computation and workflow provenance provides complementary mechanisms for documenting and validating analytical processes. Snakemake and Nextflow support structured workflow execution and dependency management (Di Tommaso et al., 2017; Köster and Rahmann, 2012), whereas CWLProv and BioCompute Objects provide standardized representations of computational provenance and analytical procedures (Alterovitz et al., 2018; Khan et al., 2019). More recently, PROV-AGENT extended provenance capture to agent interactions and decisions, while ToolGate introduced precondition- and postcondition-based checks for tool invocation and state updates (Liu et al., 2026; Souza et al., 2025). These developments provide important foundations for examining agent-directed computation. However, verifying a particular genomic analysis requires an explicit correspondence between parameters declared before execution and the arguments, parameter-file contents and execution status recorded when the underlying software is invoked. Neither the availability of domain skills nor retrospective documentation alone necessarily establishes this correspondence. Moreover, agreement between declared and executed parameters must be distinguished from the scientific validity of the selected method and the numerical correctness of its results. Evaluating these properties separately is necessary to determine which aspects of analytical reliability can be improved through domain knowledge and which require direct examination of tool execution.

Here, we investigated these questions through comparative evaluations of general-purpose agents, knowledge-driven multi-agent systems and agents equipped with domain-specific skills. To examine the correspondence between declared and executed analyses, we developed a call-level execution contract that links pre-execution parameter declarations to recorded tool invocations through cryptographic file digests, command-line argument checks and execution-state records. We implemented this approach in PigGPA, a domain-constrained agent for porcine genomic breeding that integrates 28 analytical skills spanning genomic breeding, sequence-based functional prediction and multi-omics annotation. We evaluated the systems across four representative genomic analysis tasks, comprising 84 model–system–task evaluation units with three repeated sessions per unit, together with 75 controlled mechanism checks. Numerical comparisons with HIBLUP across 14 analytical modules assessed agreement with established genetic evaluation software, while an end-to-end analysis of porcine loin muscle depth examined the application of the framework across statistical and functional genomics (Yang et al., 2021; Yin et al., 2023). By examining analytical decisions, task completion, recorded tool execution and numerical results separately, this study evaluates the contributions and limitations of domain knowledge and introduces an explicit mechanism for checking whether agent-directed computations follow their declared parameters.

## 2. Materials and Methods

### 2.1 Benchmark datasets and agent configurations

We established four representative genomic analysis tasks to evaluate the effects of domain knowledge and execution verification on autonomous scientific analysis: genome-wide association study (GWAS), heritability estimation, population structure analysis and genomic selection. The tasks used publicly available porcine genomic data from the giab048 study (Yang et al., 2021), with loin muscle depth (LMD) serving as the phenotype for analyses requiring trait information. The original dataset comprised 2,797 individuals and 258,662 single-nucleotide polymorphisms (SNPs).

Four agent configurations were evaluated: general-purpose agents without predefined domain skills, the knowledge-driven multi-agent framework BOLE, general-purpose agents equipped with domain-specific skills, and PigGPA integrating domain skills with call-level execution contracts. The general-purpose agent benchmark included Claude Code (v2.1.214) and OpenClaw (v2026.7.1-2), each evaluated using four LLMs: GLM-5.2, Qwen3.6-35B-A3B, MiniMax-M2.5 and DeepSeek-V4-Pro. Their combinations across the four analytical tasks yielded 32 evaluation units, each defined by a unique agent–model–task combination. BOLE (Wang et al., 2026) was evaluated using the same four models and four tasks, yielding 16 units. To isolate the contribution of domain skills within general-purpose agent frameworks, the PigGPA skill library was subsequently made available to Claude Code and OpenClaw under the same model and task configurations, generating another 32 units. Finally, PigGPA was evaluated on all four tasks using DeepSeek-V4-Pro, which achieved the lowest mean score among the models in the general-purpose agent benchmark. This provided a fixed, comparatively challenging model configuration for assessing the integrated system. DeepSeek-V4-Pro refers throughout to the Preview release dated 24 April 2026.

The benchmark comprised 84 evaluation units. Each unit included one reference session and three independent repeated sessions initiated with the same analytical objective and input data. All executions were isolated using Linux bubblewrap, with separate working directories to prevent exchange of intermediate files, outputs or session information between evaluation units. Comparisons between general-purpose agents with and without skills were used to assess the effects of domain knowledge under matched agent and model configurations. BOLE and PigGPA were evaluated as distinct system implementations; differences involving these systems were interpreted in the context of their respective architectures and execution procedures.

### 2.2 Evaluation criteria and reproducibility of analytical decisions

Agent performance was assessed using a standardized 100-point deduction-based scoring system comprising eight criteria. Four criteria evaluated analytical decisions: task interpretation and scope (D1-01), examination of input data (D2-01), selection of analytical methods and validation strategies (D2-02), and quality control and bias management (D2-03). Two criteria evaluated execution: agreement between planned and implemented analyses (E1-01) and completion of the execution process (E1-02). Reproducibility of analytical decisions across independent sessions was assessed separately (R-01), together with delivery of the required analytical outputs (C-01).

Each criterion was evaluated at three levels, designated L0, L1 and L2. The six decision and execution criteria and the reproducibility criterion each contributed a maximum deduction of 10 points, with deductions of 0, 5 and 10 points for L0, L1 and L2, respectively. Output delivery contributed a maximum deduction of 30 points, with corresponding deductions of 0, 15 and 30 points. An evaluation unit was classified as unsuccessful if task interpretation, execution completion or output delivery received an L2 assessment. Detailed scoring definitions and decision rules are provided in Supplementary Table S1.

Reproducibility of analytical decisions was assessed by comparing predefined methodological choices between each reference session and its three independent repetitions. The comparison included genotype quality-control strategies, statistical methods, covariate specifications and validation procedures. We recorded the consistency of individual decisions, the R-01 assessment for each evaluation unit, overall task completion and aggregate performance scores. For BOLE, consistency was assessed in the context of its predefined and ratified analytical procedures, whereas PigGPA was evaluated following autonomous workflow planning.

To further examine discrepancies between analytical plans and their implementation, execution records from the 32 skill-augmented general-purpose agent units were audited. Initial analytical plans, subsequent modifications and executed scripts were compared using predefined decision fields and parameter declarations. The audit quantified the number of modified decisions, editing events, discrepancies between declared and executed parameter values, and evaluation units in which parameters were modified before re-execution. Actual plans, scripts and execution records, rather than agent-generated final reports, served as the basis for these comparisons.

### 2.3 PigGPA architecture and domain-specific analytical skills

PigGPA was implemented as a single-agent system that separates domain knowledge from computational execution while maintaining a common environment for task planning, tool invocation and result management (Figure 1). Analytical capabilities were organized as declarative skills describing their intended applications, required inputs, permissible parameters, software dependencies and expected outputs. The LLM interpreted user objectives, examined available data and selected appropriate skills, whereas established analytical programs and scripts performed the underlying computations.

**Figure 1.**
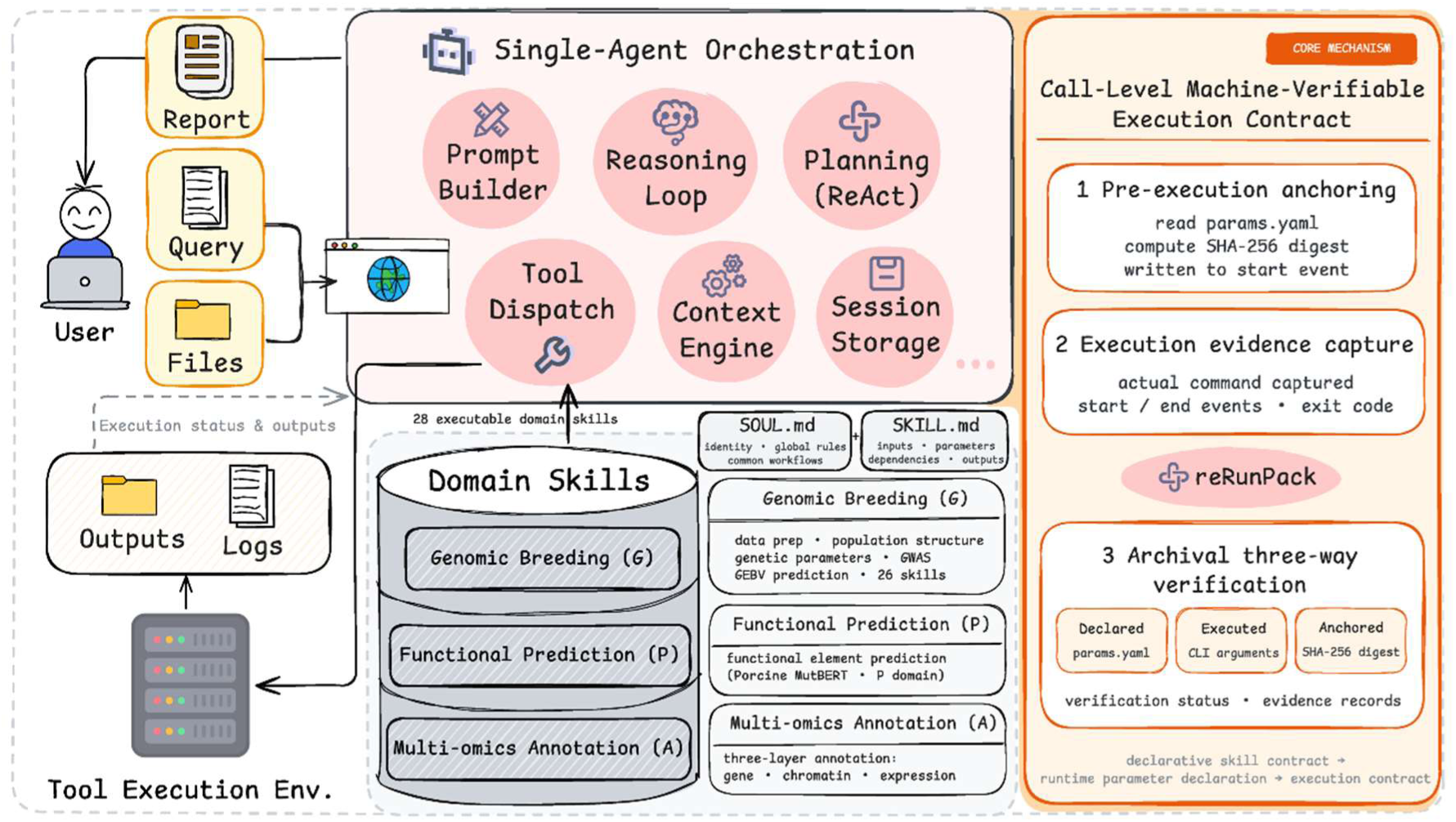
Architecture and machine-verifiable execution controls of PigGPA. The single-agent orchestration layer selects declarative domain skills and invokes analysis tools through a shared execution boundary. Before execution, the framework records the state and SHA-256 digest of params.yaml and performs a non-blocking QC-first assertion. Start and end events record commands, timestamps, exit states and parameter evidence. The verification and archive layer (reRunPack) cross-validates command-line arguments, parameter-file values and execution-time hashes, and packages scripts, dependencies, manifests and warnings for audit and replay.

The knowledge layer followed a hierarchical organization. SOUL.md defined the agent’s domain-level instructions, operating rules and available capabilities, while individual SKILL.md files specified the execution requirements of each analytical skill. Capability summaries supported initial skill selection, and detailed instructions were loaded when a skill was invoked. Users interacted with the system through either a command-line interface (CLI) or a web interface (WebUI). Analytical steps were linked through explicit input and output files, enabling results from one skill to be passed to subsequent analyses.

PigGPA incorporated 28 skills distributed across three functional domains: genomic breeding (G; 26 skills), functional element prediction (P; one skill) and genomic annotation (A; one skill). The G domain covered genotype processing and quality control, population structure analysis, GWAS, genetic parameter estimation, mixed-model analyses, genomic selection and breeding-value prediction. Computations were implemented using established software, including PLINK, GCTA and ADMIXTURE, together with existing statistical procedures (Alexander et al., 2009; Chang et al., 2015; Purcell et al., 2007; Yang et al., 2011). The complete skill inventory is provided in Table 1.

**Table 1.** Catalogue of PigGPA analytical skills.

| Skill identifier | Main function | Method or software |
| --- | --- | --- |
| **G \| Genomic breeding — Data preparation** |  |  |
| genotype-import* | Import VCF or PLINK binary data | bcftools; PLINK |
| genotype-qc* | Genotype quality control based on MAF, HWE and missingness | PLINK 1.9 |
| qc | Integrated diagnostics for genotype, phenotype and metadata, including PCA, IBS and NJ trees | PLINK; Python |
| internal-impute | Reference-panel-free imputation using multiple genotyping arrays and mutual population references | Beagle 5.x; PLINK |
| **G \| Genomic breeding — Association analysis** |  |  |
| plink-gwas* | Genome-wide association analysis | PLINK linear or logistic regression |
| gcta-gwas* | Mixed-model association analysis | GCTA-MLMA |
| **G \| Genomic breeding — Population structure** |  |  |
| pca | Principal component analysis | Patterson PCA |
| admixture* | Estimate ancestry proportions | ADMIXTURE |
| ld-pruning* | Linkage disequilibrium pruning | PLINK LD pruning |
| ld | Estimate linkage disequilibrium and its decay | PopLDdecay; Hill–Weir model |
| ld-score | Estimate LD scores | LDSC |
| allele-genotype-frequency | Estimate allele and genotype frequencies and test HWE | PLINK; HWE tests |
| nj-tree | Construct NJ trees from IBS distance matrices with circular or rectangular layouts | Biopython Phylo |
| **G \| Genomic breeding — Genetic parameters** |  |  |
| homozygosity | Assess homozygosity | PLINK |
| inbreeding | Estimate inbreeding from GRM diagonals, homozygosity proportions or excess homozygosity | PLINK; GCTA |
| roh | Detect ROH and estimate F<sub>ROH</sub>, with breed-level bar plots | PLINK --homozyg |
| relationship-matrix | Construct PRM, GRM and HRM | VanRaden or Yang GRM; PRM; HRM ($0.95 \times G_{adj} + 0.05 \times A$) |
| snp-effect | Estimate SNP effects | RR-BLUP |
| heritability* | Estimate heritability using GREML and ICC | GCTA-GREML; ICC |
| **G \| Genomic breeding — Breeding models** |  |  |
| single-trait | Single-trait evaluation using SSGBLUP | SSGBLUP (AI-REML, EM-REML or HE) |
| multi-trait | Multi-trait evaluation using multivariate mixed models | Multivariate mixed model with GRM eigendecomposition |
| repeated-records | Repeated-records analysis with permanent environmental effects | Mixed model with permanent environmental effects; Woodbury identity |
| gxe | Model genotype-by-environment interactions | Mixed model with a GRM Hadamard-product interaction matrix |
| linear-mixed | Fit LM (HE), PBLUP, GBLUP and SSGBLUP models | HE regression; Henderson equations |
| **G \| Genomic breeding — Prediction applications** |  |  |
| gebv-gprs-prediction | Predict GEBV and evaluate GPRS | $GEBV = X \cdot \hat{\beta}$; GPRS evaluation |
| genomic-selection* | Cross-validation of six genomic prediction models | BGLR/rrBLUP: BayesA, BayesB, BayesC, BRR, BL and GBLUP |
| **P \| Functional element prediction** |  |  |
| pig-mutbert | Predict porcine regulatory elements from genomic sequences across seven tasks and multiple tissues | Porcine MutBERT |
| **A \| Genomic annotation** |  |  |
| pig-annotation-track | Integrate gene models, chromatin states and tissue expression in three-layer tracks | Ensembl gene models; chromatin-state segmentation; tissue expression matrices |

G, genomic breeding (26 skills); P, functional element prediction (one skill); A, genomic annotation (one skill). Each skill specifies its inputs, parameters, dependencies and outputs; computations reuse established software and statistical routines. \* Skills inherited from the BOLE knowledge base (Wang et al., 2026). MAF, minor allele frequency; HWE, Hardy–Weinberg equilibrium; IBS, identity by state; NJ, neighbour joining; LD, linkage disequilibrium; ROH, runs of homozygosity; F\_ROH, ROH-based inbreeding coefficient; PRM, pedigree relationship matrix; GRM, genomic relationship matrix; HRM, hybrid relationship matrix; GEBV, genomic estimated breeding value.

The P domain incorporated the previously developed Porcine MutBERT family of genomic foundation models (Long et al., 2026), supporting sequence-based prediction of seven functional features: ATAC accessibility, CTCF binding, enhancers, promoters, H3K27ac, H3K4me1 and H3K27me3. The skill accepted DNA sequences, FASTA files, CSV files containing sequence information, and genomic positions or intervals. It automated reference sequence extraction, model selection, inference and output processing. Model pretraining and the original predictive performance evaluations have been reported previously; the present study focused on standardized inference and integration with other analytical operations.

The A domain generated integrated genomic annotation tracks from SNP positions, genomic intervals or gene identifiers using the Sscrofa11.1 reference assembly. The resulting visualizations combined Ensembl gene models, ChromHMM annotations comprising 15 chromatin states across 14 tissues, and expression profiles from 34 tissues or cell populations (Ernst and Kellis, 2017, 2012; Pan et al., 2021; Teng et al., 2024).

Cross-domain workflows were assembled through genomic coordinates, sample identifiers and intermediate data files. For example, SNPs identified by GWAS were converted into reference genomic sequences before being submitted to Porcine MutBERT. The corresponding genomic positions were subsequently used to retrieve functional annotations and generate visualization tracks. This organization enabled heterogeneous analytical tools to operate within a single workflow while retaining the records associated with each execution step.

### 2.4 Call-level execution contracts and verification procedures

We developed a call-level execution contract mechanism to examine whether recorded tool invocations agreed with parameters declared before execution. The mechanism operated at the boundary between agent planning and computational execution, independently of the agent’s subsequent natural-language description of the analysis.

Before a tool invocation, the system inspected the corresponding parameter file (params.yaml) and, when available, recorded its declared values and SHA-256 digest together with the command and working directory. The digest provided a reference for detecting changes in the recorded parameter-file contents. For skills involving statistical inference, the system additionally checked for the presence of expected quality-control parameter files. Missing quality-control evidence generated a non-blocking QC-first warning without automatically modifying the agent’s analytical decisions.

Each invocation generated start and end records containing a common invocation identifier and execution-state information, including the termination status and exit code. During subsequent archiving, recorded command-line arguments were compared with declared parameter values, and parameter-file digests were compared across the execution interval. The system also examined whether parameter modifications occurring during execution were accompanied by corresponding records. Invocation identifiers were used to associate start and end events and identify incomplete or abnormally terminated tool calls.

Execution records, parameter files, logs and analytical outputs were organized into a reproducibility package termed reRunPack. Verification outcomes were recorded separately from task completion. Missing parameter declarations, inconsistent values and incomplete execution events were retained as distinct evidence states rather than automatically classified as successful verification.

The mechanism was designed to examine agreement between declared parameters and observable execution records. It did not enforce all quality-control requirements, inspect every operation performed internally by the underlying software, or independently establish the scientific validity of the selected analytical methods. Its verification scope was therefore determined by the parameter declarations and execution events available for each invocation.

### 2.5 Controlled verification tests and numerical benchmarking

The execution-contract mechanism was evaluated through 75 controlled checks comprising parameter anchoring (12 checks), archive verification (34 checks) and end-to-end integration (29 checks). Test conditions included missing parameter files, disagreement between declared and command-line parameter values, unrecorded modifications to parameter files during execution, and invocation records lacking corresponding termination events. Expected verification outcomes were specified for each condition. Normal invocations were also examined to assess whether the mechanism incorrectly reported discrepancies.

Numerical consistency was evaluated by comparing PigGPA with HIBLUP (Yin et al., 2023) across 14 shared analytical modules. A simulated dataset containing 10,000 individuals, 100,000 SNPs and 18 autosomes was used. The principal comparisons employed the first 1,000 individuals and 5,556 SNPs on chromosome 1, while genomic prediction was evaluated using separate training and validation subsets of 1,000 individuals each. Agreement was assessed using Pearson correlation coefficients, mean squared errors, maximum absolute differences and comparisons of estimated variance components and model convergence. Differences arising from distinct statistical estimators, parameterizations or numerical implementations were examined separately.

Additional implementation tests comprised 100 constructed natural-language queries, 26 executable workflows covering the G-domain skills, and five abnormal-input scenarios. Queries represented eight analytical categories and three difficulty levels. Abnormal-input tests included missing files, invalid minor allele frequency thresholds, ambiguous analytical requests, nonexistent file paths and incompatible parameters. These tests assessed intent routing, implementation coverage and error handling, independently of the multistep agent benchmark.

To examine the portability of archived analyses, two reRunPack instances were transferred to separate working directories and re-executed. Their completion status and selected output artifacts were compared against predefined consistency criteria.

### 2.6 End-to-end genomic analysis and functional interpretation

An end-to-end application was conducted using the publicly available LMD dataset to examine whether PigGPA could coordinate statistical genetics, sequence-based functional prediction and multi-omics annotation within a single analytical workflow (Yang et al., 2021). Starting from 2,797 individuals and 258,662 SNPs, the workflow included input inspection, genotype quality control, alignment of genotype, phenotype and covariate records, mixed-model GWAS, candidate-region sequence extraction, Porcine MutBERT inference and integrated genomic annotation.

Workflow completion was evaluated according to the generation of essential intermediate and final artifacts and their successful use by downstream operations. GWAS results were compared with the original study using genome-wide association patterns, genomic inflation factors and association statistics for shared SNPs in the SSC7:30–38 Mb region. Pearson and Spearman correlations were used to assess agreement in regional association signals.

Functional predictions were generated for selected lead SNPs across available task–tissue combinations. Predicted regulatory features were examined together with genomic annotations, chromatin states and expression profiles. These analyses were intended to demonstrate the integration and traceability of computational evidence, rather than experimentally establish the regulatory activity or causal effects of candidate variants.

## 3. Results

### 3.1 General-purpose agents exhibit substantial variation in analytical decisions

We first evaluated general-purpose agents performing genomic analyses without predefined domain-specific skills. Across four analytical tasks and four underlying language models, Claude Code completed 15 of 16 evaluation units, achieving a mean score of 77.50/100, whereas OpenClaw completed 13 of 16 units with a mean score of 69.38/100. Together, the two frameworks completed 28 of 32 units, with an overall mean score of 73.44/100. Model-level mean scores ranged from 70.63 to 79.38, with GLM-5.2 achieving the highest score and DeepSeek-V4-Pro the lowest (Figure 2).

**Figure 2.**
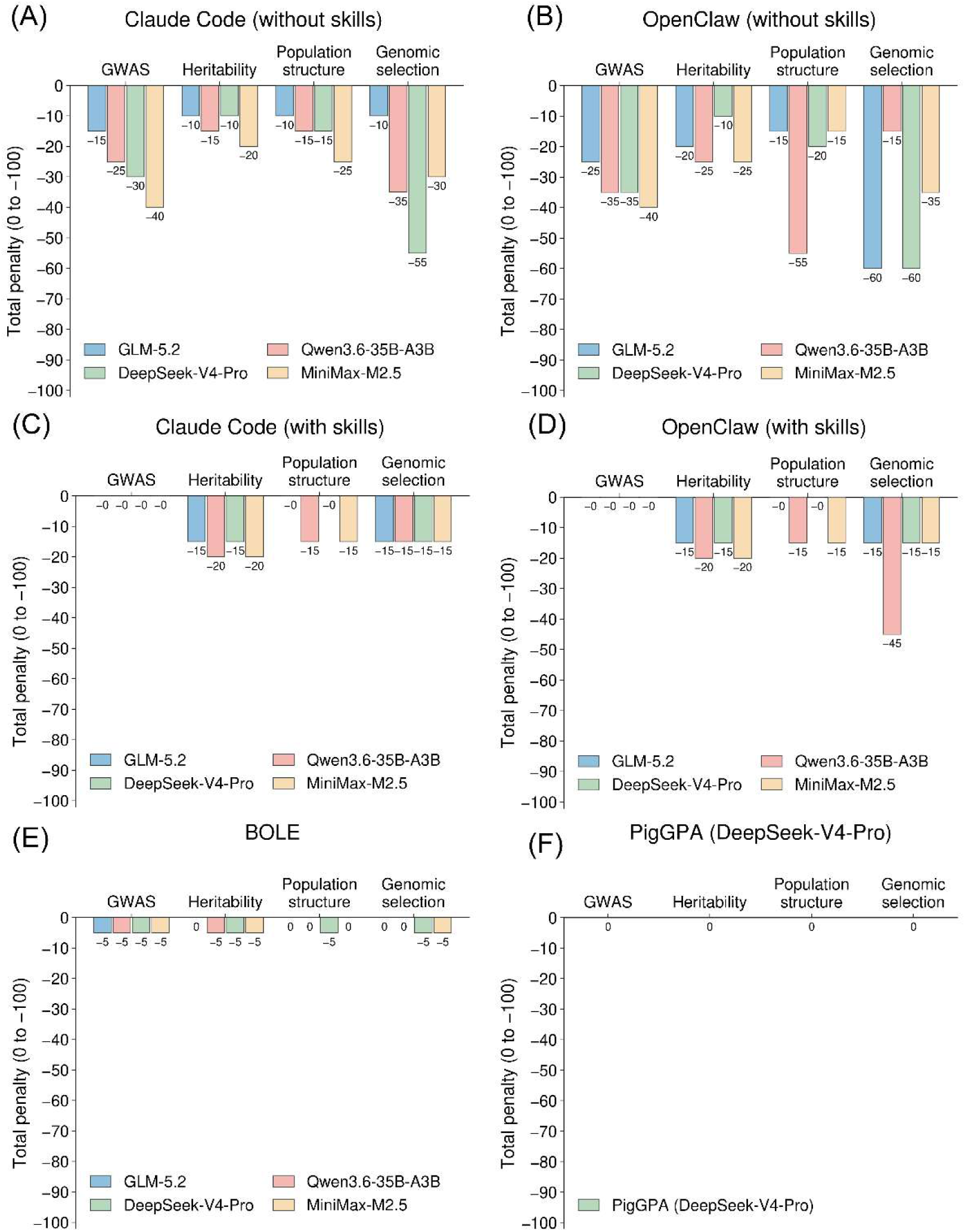
Total score deductions across benchmark tasks and model configurations. (a) Claude Code; (b) OpenClaw; (c) BOLE; (d) PigGPA under the DeepSeek-V4-Pro stress test. Bar values represent the total deduction from the 100-point baseline for each model × task unit and do not decompose deductions by scoring block. PigGPA results therefore describe only the tested model configuration.

Despite completing most tasks, both frameworks showed substantial variation in analytical decisions across independent sessions. All 32 evaluation units received deductions under the decision reproducibility criterion (R-01). The pooled consistency of predefined analytical decisions across three repeated sessions was 58.8% for Claude Code and 57.5% for OpenClaw. Disagreements primarily involved genotype quality-control strategies, covariate selection, evaluation metrics and combinations of statistical methods, whereas some individual threshold parameters were comparatively stable.

Failures were concentrated in tasks requiring extended computation or coordination of multiple analytical steps. All four unsuccessful units received the maximum deduction for execution completion, including three genomic selection tasks. Recorded failures included sessions terminating before background computations had finished, repeated attempts to repair failing analyses until the available context was exhausted, and tool or host-process failures. All three unsuccessful OpenClaw units were associated with context truncation. Thus, although general-purpose agents frequently generated the required analytical outputs, successful completion did not imply consistent methodological decisions across repeated sessions.

### 3.2 Domain knowledge improves analytical performance but does not ensure agreement between planned and executed analyses

We next examined whether structured domain knowledge could reduce the variability observed in general-purpose agents. The knowledge-driven multi-agent framework BOLE completed all 16 evaluation units, achieving a mean score of 96.88/100. Scores across the four underlying models ranged from 95.00 to 98.75, compared with 70.63–79.38 for the general-purpose agents (Figure 2). Under the DeepSeek-V4-Pro configuration, all four BOLE tasks received no deduction for decision reproducibility, whereas the corresponding general-purpose agent configurations showed varying degrees of inconsistency.

BOLE also completed all four genomic selection evaluations, each involving comparisons among six prediction models. However, the computational settings were not identical across its model configurations: three used 197,702 SNPs and 10-fold cross-validation, whereas the GLM-5.2 configuration used 874 LD-pruned SNPs and fivefold cross-validation because of memory limitations. These findings indicate that BOLE achieved high task completion and stable analytical plans under the tested conditions, although differences from general-purpose agents cannot be attributed exclusively to domain knowledge because their workflow organization and decision procedures also differed.

To assess the contribution of reusable domain skills more directly, we equipped Claude Code and OpenClaw with the PigGPA skill library while retaining the same four models and analytical tasks. Task completion increased from 28/32 to 31/32 units, and the mean score increased from 73.44 to 89.06/100. Claude Code achieved 90.00/100 and OpenClaw 88.13/100. All 32 skill-augmented units received no deductions for input-data examination (D2-01) or method and validation design (D2-02), indicating improvements in these aspects of analytical planning. The number of units without the required delivered outputs decreased from four to one. In the remaining unsuccessful unit, a genomic selection analysis running under OpenClaw with Qwen3.6-35B-A3B generated the principal prediction results but saved them outside the expected delivery directory (Figure 2).

These improvements did not eliminate discrepancies between analytical plans and their implementation. Although genotype quality control was included in the planned analyses for all 32 skill-augmented units, execution records showed no genotype-filtering operation in 20 units (62.5%). Auditing the planning and execution records further identified modifications to 15 of 192 predefined decision fields (7.81%) after the initial plans had been generated, comprising 40 editing events. Of 1,377 parameter declarations, 50 (3.63%) disagreed with the corresponding executed scripts. At least three of the 32 units (9.38%) modified parameters and reran analyses during execution; including reruns associated with infrastructure timeouts increased this number to six.

These observations distinguish the benefits of providing established analytical methods from the separate requirement to verify their implementation. Domain knowledge and reusable skills improved task completion and several aspects of analytical planning, but discrepancies between declared procedures and executed operations remained detectable in the skill-augmented agents.

### 3.3 PigGPA combines task completion with auditable execution

We integrated the domain-specific analytical skills and execution-contract mechanism into PigGPA and evaluated the resulting system using DeepSeek-V4-Pro, the lowest-scoring model in the general-purpose agent benchmark. PigGPA completed all four genomic breeding tasks and achieved 100/100 under the standardized scoring criteria for each task (Figure 2). Across three independent repetitions of each task, all predefined key analytical decisions agreed with the corresponding reference sessions, yielding 12/12 consistent repeated sessions.

Execution auditing complemented task-level performance by linking analytical actions to inspectable records. The reference workflows followed their planned analytical stages, and the audited settings agreed with recorded execution arguments or documented skill defaults where comparable evidence was available. All 75 checks in the implementation acceptance record met their expected outcomes, supporting the recording and archival mechanisms under the tested conditions. Together, these findings demonstrate the feasibility of combining successful task completion with traceable and auditable execution.

We independently assessed the numerical results of PigGPA against HIBLUP across 14 shared analytical modules (Figure 3). Homozygosity, heterozygosity and pedigree relationship matrix estimates were highly concordant, with corresponding Pearson correlations of 1.0000. Genomic and hybrid relationship matrices were also strongly correlated between implementations (r = 0.9990 and 0.9991, respectively). The reported GBLUP predictive correlations differed by 0.000247.

**Figure 3.**
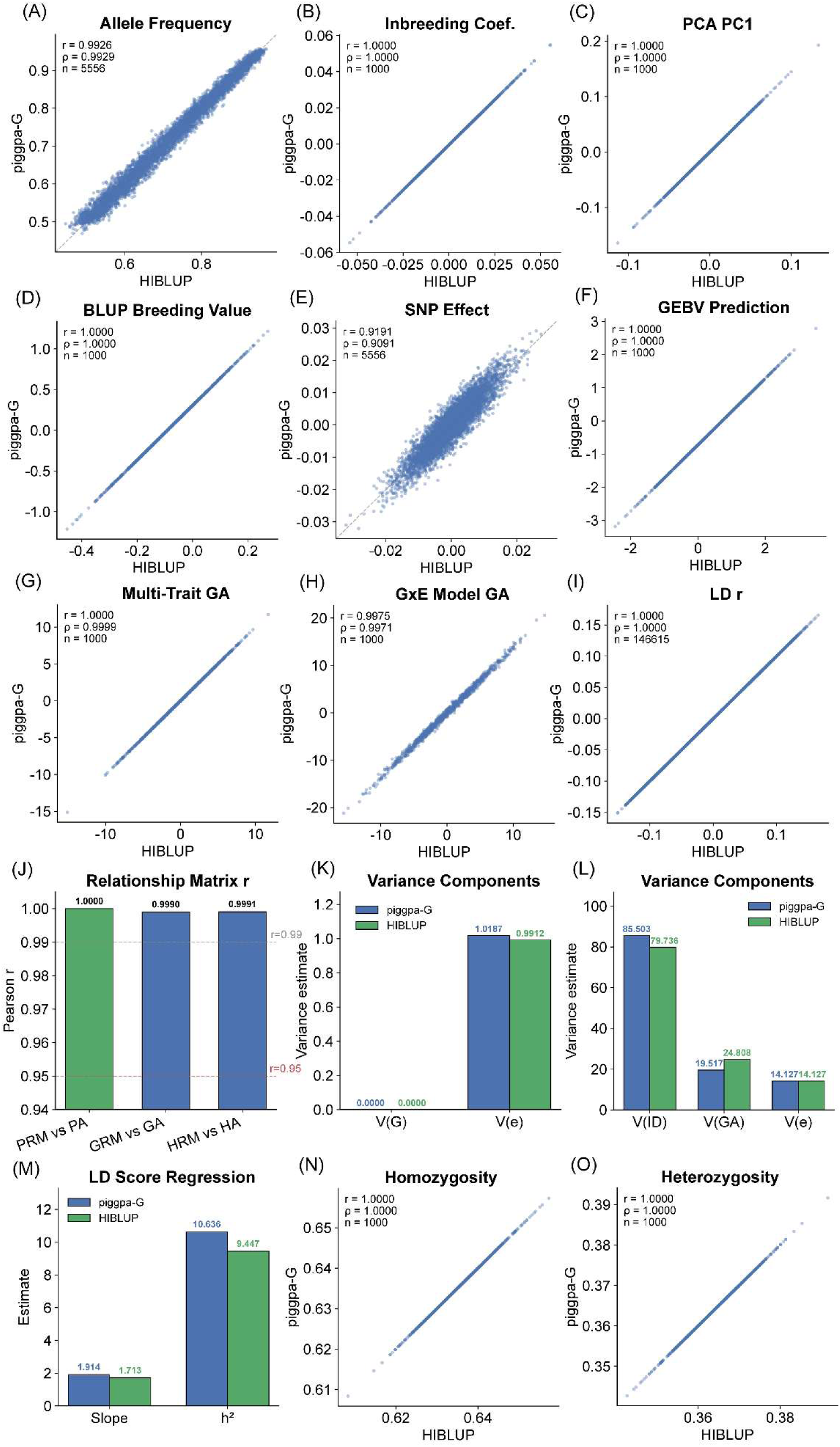
Numerical comparison between PigGPA and HIBLUP (v1.6.0) across 14 shared modules, using simulated pig population data (1,000 individuals × 5,556 SNPs on chr1) under identical input conditions. Scatter plots (a–i, n, o) show element-wise agreement between PigGPA (y axis) and HIBLUP (x axis), annotated with Pearson r and Spearman ρ; bar plots (j–m) summarize module-level statistics. (a) Allele frequencies (r = 0.9926, n = 5,556 SNPs; A1 frequencies aligned between tools). (b) Inbreeding coefficients (r = 1.0000, n = 1,000). (c) First principal component (r = 1.0000, n = 1,000). (d) BLUP breeding values (r = 1.0000, n = 1,000; both tools use the genomic relationship matrix GA). (e) SNP effects (r = 0.9191, n = 5,556; PigGPA RR-BLUP versus HIBLUP GBLUP). (f) GEBV predictions (r = 1.0000, n = 1,000). (g) Multi-trait model additive effects GA (r = 1.0000, n = 1,000). (h) G×E model GA (r = 0.9975, n = 1,000). (i) Pairwise LD Pearson r (r = 1.0000, n = 146,615 SNP pairs; both tools report Pearson r rather than r²). (j) Relationship-matrix correlations: PRM vs PA (r = 1.0000), GRM vs GA (r = 0.9990), HRM vs HA (r = 0.9991); dashed lines mark r = 0.99 and r = 0.95. (k) Single-trait variance components (AI-REML): V(G) ≈ 0 in both; V(e) = 1.019 (PigGPA) vs 0.991 (HIBLUP). (l) Repeated-records variance components: V(ID) = 85.50 vs 79.74, V(GA) = 19.52 vs 24.81, V(e) = 14.13 in both; total variance differs by 0.4%, and the V(ID)/V(GA) allocation difference is attributable to GRM scaling (VanRaden vs Su et al.). (m) LD-score regression: slope = 1.914 (PigGPA) vs 1.713 (HIBLUP; ∼11% difference between OLS and two-step estimators); h² = 10.636 vs 9.447, both exceeding 1 as expected for strong-signal in-silico data at N = 1,000. (n) Homozygosity (r = 1.0000, n = 1,000; maximum absolute difference ∼6 × 10⁻⁷). (o) Heterozygosity (r = 1.0000, n = 1,000). Non-converged or biologically inadmissible estimates are retained as diagnostic results and are not treated as evidence of equivalence.

Some more complex analyses showed differences between implementations. In the repeated-records model, PigGPA converged after nine iterations, whereas HIBLUP had not converged within its specified limit of 20 iterations; nevertheless, both yielded a residual variance of approximately 14.1266. LD Score regression produced heritability estimates of 10.6357 and 9.4469 for PigGPA and HIBLUP, respectively. Because both estimates exceeded the conventional range of heritability, they were used only to characterize differences between implementations under the simulated conditions and were not interpreted biologically. Overall, the numerical comparisons supported close agreement for several core genetic analyses, while revealing estimator-dependent differences that require separate interpretation.

To assess the ability to rerun archived analyses, we transferred two reRunPack instances to independent working directories. Both were successfully re-executed, and their selected output artifacts met the predefined consistency criteria. Additional implementation tests showed that all 26 G-domain executable workflows completed successfully, all 100 constructed queries were routed to their intended skills, and all five abnormal-input scenarios produced the expected responses. These findings separately support implementation coverage, numerical consistency and limited workflow replay capability, rather than providing interchangeable measures of analytical correctness or execution verification.

### 3.4 PigGPA integrates genetic association, sequence-based prediction and multi-omics annotation

Finally, we evaluated PigGPA in an end-to-end analysis of porcine loin muscle depth, linking statistical genetics with sequence-based functional prediction and multi-omics annotation. Starting from 2,797 individuals and 258,662 SNPs, the system performed input examination and genotype quality control, retaining 2,770 individuals and 197,702 SNPs. Subsequent operations included genotype–phenotype matching, mixed-model GWAS, extraction of reference sequences for candidate variants, Porcine MutBERT inference and integrated genomic annotation (Figure 4).

**Figure 4.**
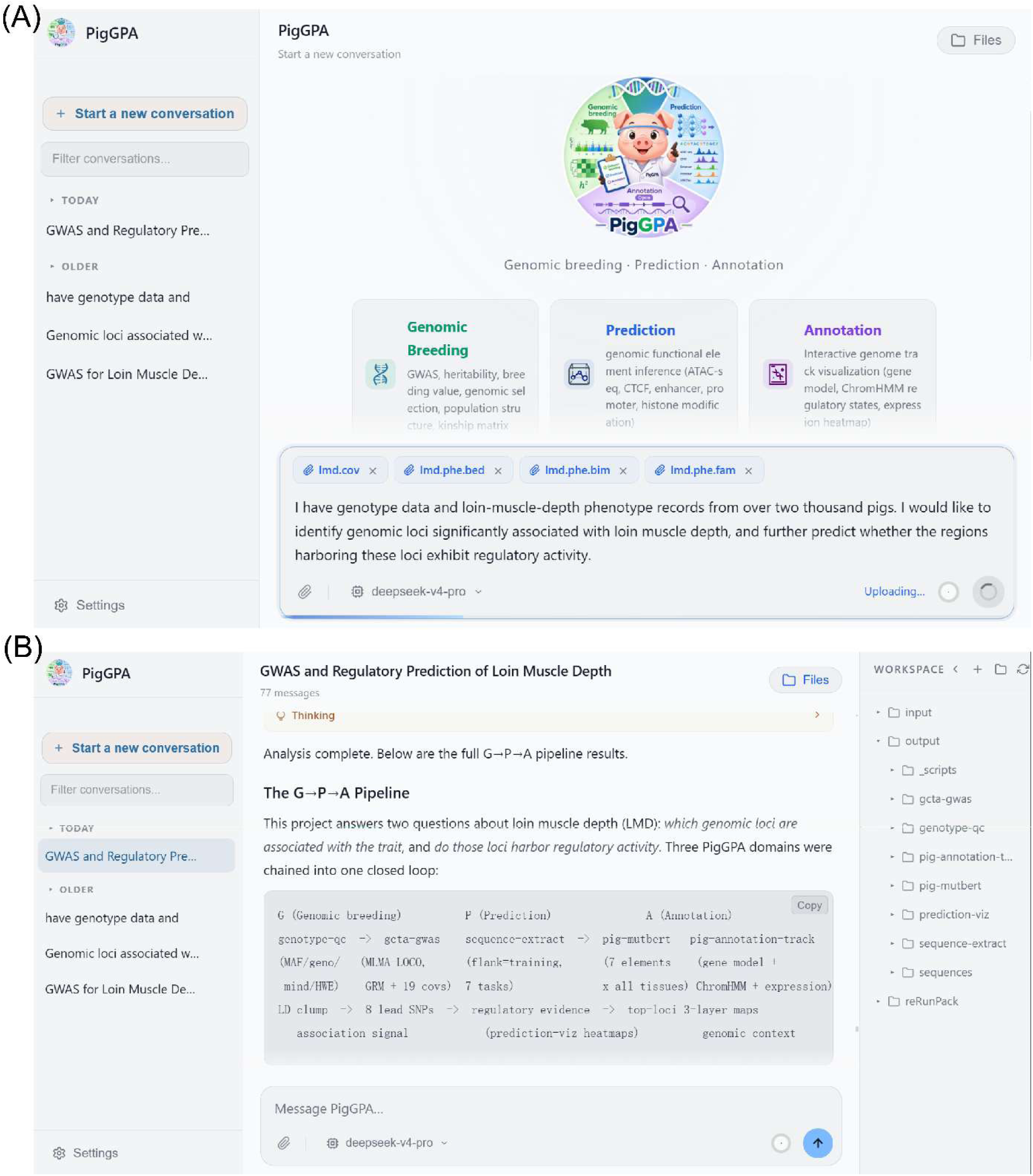
PigGPA interface during the LMD case study. (a) Conversation setup: the user states the goal of “identifying genomic loci significantly associated with loin muscle depth and predicting their regulatory activity” in a natural-language prompt and attaches the genotype and covariate files (lmd.phe.bed/.bim/.fam, lmd.cov); the panel also shows the skill scope covered by the three domains, G (Genomic Breeding), P (Prediction) and A (Annotation). (b) Execution and workspace view: the conversation list on the left, the conversation area in the middle, and the WORKSPACE panel on the right listing the files actually written to disk under input/, output/ and each domain’s skill directories, so that each step’s outputs can be checked directly.

The association patterns showed broad agreement with the published study (Figure 5A). Within the SSC7:30–38 Mb region, −log₁₀(P) values for 289 shared variants were strongly correlated (Pearson r = 0.9512; Figure 5B), indicating concordance on the same source dataset. Beyond this comparison, the MLMA-LOCO workflow identified 24 SNPs at P < 5 × 10⁻⁸, which LD clumping reduced to eight candidate lead SNPs, four each on SSC7 and SSC13. The strongest signals were SSC7:36,571,258 (P = 6.29 × 10⁻¹¹) and SSC13:82,021,256 (P = 4.99 × 10⁻¹⁰). These candidates were carried forward for sequence-based functional prediction and multi-omics annotation.

**Figure 5.**
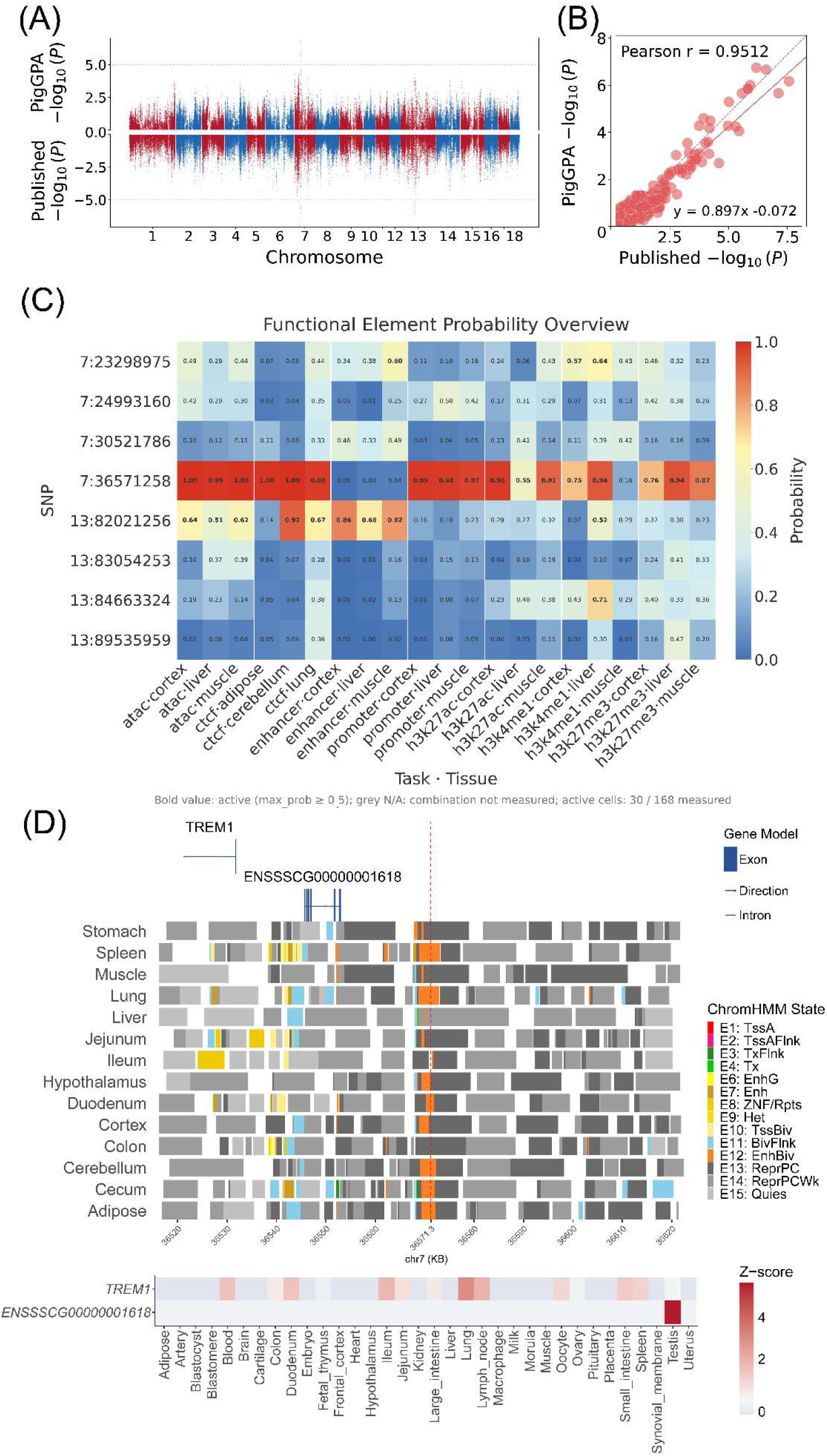
Comparison of association patterns and functional annotation of candidate variants for porcine loin muscle depth. (A) Mirror Manhattan plot comparing PigGPA association results (top) with the published results (bottom, with the ordinate inverted); alternating colours distinguish chromosomes. (B) Comparison of −log₁₀(P) values for 289 shared variants in the SSC7:30–38 Mb region (Pearson r = 0.9512). The solid red line shows the fitted regression (y = 0.897x − 0.072), and the dashed grey line denotes y = x. Panels A and B assess concordance on the same source dataset and do not constitute independent replication. (C) Porcine MutBERT prediction probabilities for eight candidate lead SNPs across 21 task–tissue combinations. Rows indicate SNPs ordered by genomic position; columns represent seven prediction tasks with three compatible tissues per task. Cell values are the maximum predicted probabilities, and bold values indicate probabilities ≥ 0.5 (30 of 168 combinations). (D) Integrated annotation of the region surrounding SSC7:36,571,258 (±50 kb), showing gene models, 15-state ChromHMM annotations across 14 tissues and expression Z-scores across 34 tissues or cell populations. The red dashed line marks the SNP position. Predictions and annotations provide computational context for candidate prioritization and do not establish regulatory function or causality.

The eight candidate lead SNPs were evaluated using Porcine MutBERT across seven prediction tasks, each involving three compatible tissues. The analysis covered 21 task–tissue combinations across six distinct tissues, yielding 168 SNP–task–tissue predictions. Thirty combinations had a maximum predicted probability of at least 0.5 (Figure 5C). Input sequence windows ranged from 400 to 1,000 bp according to the task-specific inference settings.

Among the evaluated variants, SSC7:36,571,258 received high predicted ATAC accessibility probabilities in muscle, cortex and liver (1.00, 1.00 and 0.99, respectively), with positive prediction labels in 17 of the 21 task–tissue combinations. Another candidate, SSC13:82,021,256, showed predicted features associated primarily with CTCF binding and enhancer activity (Figure 5C). ChromHMM annotations assigned SSC7:36,571,258 to state E12 (EnhBiv) in all six tissues examined in the prediction analysis, placing this candidate in a putative regulatory context (Figure 5D). These findings represent sequence-model predictions and existing genomic annotations rather than direct experimental measurements of regulatory activity.

The annotation workflow further aligned candidate variants with gene structures, tissue-specific chromatin states and expression profiles to generate integrated genomic tracks (Figure 5D). Input files, parameter declarations, recorded tool invocations and output artifacts remained linked across the successive analytical stages. This application demonstrates that PigGPA can coordinate heterogeneous tools and retain inspectable execution records from genetic association analysis through functional interpretation. The computational predictions provide additional context for candidate variants but do not establish their causal roles or underlying regulatory mechanisms.

## 4. Discussion

### 4.1 Domain knowledge improves analytical performance but does not ensure consistent execution

Our results highlight an important distinction between selecting appropriate analytical methods and executing them as specified. General-purpose agents completed most genomic analysis tasks but showed substantial variation in methodological decisions across independent sessions. Incorporating structured domain knowledge improved analytical performance, as demonstrated by the high task completion and relatively consistent performance of BOLE across different language models. A more direct comparison, in which the same general-purpose agents were evaluated with and without domain-specific skills, further showed that skill availability increased task completion from 28/32 to 31/32 and improved the mean evaluation score from 73.44 to 89.06. These findings support the value of providing scientific agents with established analytical procedures, explicit software interfaces and domain-specific methodological guidance.

However, improvements in analytical planning did not necessarily translate into consistent implementation. Despite the availability of domain skills, 20 of the 32 skill-augmented evaluations did not execute genotype filtering that had been included in their analytical plans, and 50 of 1,377 declared parameters differed from those in the corresponding execution scripts. These discrepancies suggest that access to validated analytical procedures does not, by itself, establish whether the intended procedures were followed. Importantly, modifications to an analytical plan are not inherently erroneous: changes may be justified by data characteristics, computational constraints or intermediate results. The critical requirement is that such changes are documented and that the resulting computation can be compared with the parameters ultimately declared for execution. This distinction is particularly relevant to scientific agents, whose ability to adapt analytical workflows is useful but can complicate the reconstruction of how a result was obtained.

The comparison with BOLE also requires consideration of differences in system design. BOLE uses structured workflows and plan adjudication to constrain analytical decisions, whereas the skill-augmented general-purpose agents retain greater flexibility in planning and executing tasks. Consequently, their performance differences cannot be attributed solely to domain knowledge. Nevertheless, the combined findings indicate that domain knowledge and reusable analytical skills address only part of the problem. They can improve method selection and task completion, but verification of actual tool execution requires additional information beyond the agent’s plan and final report.

### 4.2 Call-level execution contracts enable direct examination of declared and recorded computations

To address this limitation, we introduced call-level execution contracts that associate parameters declared before execution with the corresponding tool invocation records. The contribution of this approach is not the use of cryptographic hashing or execution logging individually, but their integration into a verification procedure that checks parameter declarations, recorded command-line arguments, parameter-file changes and invocation termination states. By evaluating these records independently of the agent’s final description, the system can distinguish agreement between declared and recorded execution from detectable discrepancies and cases in which the available evidence is insufficient.

Controlled evaluations supported the ability of this mechanism to identify the tested discrepancies. All 75 predefined checks produced their expected outcomes, including the detection of four representative abnormal conditions involving missing parameter files, inconsistent command-line values, unrecorded parameter changes and incomplete invocation records. No false-positive warnings were observed among 10 normal invocations. In the PigGPA benchmark, all 34 scalar parameters for which comparable declarations and execution records were available agreed with their pre-execution values. These findings establish the feasibility of checking specified aspects of agent-directed computation through structured execution records rather than relying exclusively on retrospective descriptions.

An important observation was that successful execution and complete verification evidence were not equivalent. Among 22 analytical skill invocations requiring parameter declarations, only 10 had corresponding parameter files available for comparison; the remaining 12 were explicitly recorded as having incomplete parameter evidence. This distinction is essential because treating missing evidence as successful verification would obscure the limitations of the records being examined. The current results therefore demonstrate both the utility of call-level verification and the need to improve its coverage. Moreover, the controlled tests included a limited number of deliberately introduced abnormalities and normal invocations, and do not establish detection performance across all possible execution failures.

Execution contracts complement rather than replace established approaches to reproducible computation. Workflow systems such as Snakemake and Nextflow provide dependency management, structured execution and workflow rerunning (Di Tommaso et al., 2017; Köster and Rahmann, 2012), while CWLProv and BioCompute Objects provide representations of computational procedures and provenance (Alterovitz et al., 2018; Khan et al., 2019). More recent approaches have extended provenance and verification to agent interactions and tool use (Liu et al., 2026; Souza et al., 2025). Our implementation addresses a narrower requirement arising when agents select tools and specify parameters during runtime: establishing an explicit correspondence between pre-execution declarations and the observable records of individual analytical invocations. This verification layer can operate alongside existing workflow managers and provenance systems without requiring their replacement.

The resulting checks also have a defined scope. Agreement between declared and recorded command-line parameters does not prove that a statistical model was correctly specified, that its assumptions were satisfied or that the underlying software produced accurate results. Similarly, an SHA-256 digest can detect differences between recorded file states but does not independently establish which values were consumed internally by a program. The distinction between execution verification and numerical correctness is reflected in our separate comparisons with HIBLUP, which showed close agreement for several core quantitative-genetic calculations but also revealed differences in more complex estimators. These evaluations address complementary questions and should not be interpreted as interchangeable evidence of analytical reliability.

### 4.3 Limitations and broader implications

Several limitations define the scope of the present findings. First, the execution contracts cover parameters and events exposed through the implemented recording interfaces. Settings supplied through environment variables, software defaults, internal script assignments or external configuration files may affect computational results without appearing in the captured declarations. Changes in reference genomes, annotation databases, software versions or random-number generation can also alter results even when recorded command-line arguments remain unchanged. More comprehensive verification will therefore require capturing additional execution dependencies and extending parameter checks beyond the current interfaces. Furthermore, cryptographic digests do not by themselves provide tamper-resistant provenance when the components generating and storing execution records are subject to modification. Stronger guarantees would require an independently controlled recording or validation mechanism.

Second, the present evaluation establishes reproducibility and execution agreement only under the tested configurations. PigGPA was evaluated with a single underlying language model across four representative genomic breeding tasks, limiting conclusions about the effects of model choice or task complexity. Although its repeated sessions showed consistent predefined analytical decisions, this does not establish that all possible analytical choices would remain unchanged under different data conditions. Similarly, successful re-execution of two archived analyses demonstrates the feasibility of workflow replay in the tested environments, but not complete reproducibility across software versions or computing infrastructures. The remaining gaps in parameter-file coverage further emphasize that verification completeness must be evaluated separately from task completion.

Third, the genomic application demonstrates workflow integration rather than the discovery of experimentally validated regulatory mechanisms. PigGPA connected association analysis with sequence-based functional prediction and multi-omics annotation, while retaining records linking successive analytical steps. The association results showed regional agreement with the published analysis of the same underlying dataset, but this comparison does not constitute independent replication. The elevated genomic inflation factor observed in the current LMD analysis also requires further investigation before the significant associations can be considered robust. Likewise, Porcine MutBERT predictions and chromatin annotations provide computational evidence for candidate regulatory regions but do not establish their functional or causal effects. These distinctions are important because complete execution records cannot compensate for limitations in the underlying statistical analysis or biological evidence.

Although PigGPA was developed and evaluated in porcine genomics, the execution-contract mechanism itself depends on tool interfaces and parameter records rather than species-specific biological knowledge. Its underlying design could therefore be adapted to other computational research settings in which agents dynamically select analytical methods and invoke external programs. Such applications would require appropriate domain skills, explicit parameter specifications and verification rules suited to the underlying tools. Demonstrating this transferability will require independent evaluations across other analytical systems and scientific domains.

Taken together, our findings show that domain knowledge and execution verification address different requirements of autonomous scientific analysis. Domain-specific skills improve the ability of agents to select and complete specialized computational procedures, whereas call-level execution contracts provide a means of examining whether recorded computations agree with their declared parameters. Their combination does not eliminate the need for methodological assessment, numerical validation or independent scientific verification. It does, however, make an important part of the computational process directly inspectable, providing a practical basis for evaluating and improving agent-directed scientific workflows.

## Data and code availability

The porcine genomic data analyzed in this study were obtained from the publicly available dataset reported by Yang et al. (2021) (DOI: 10.5524/100894). The Porcine MutBERT models and associated resources were described previously by Long et al. (2026). The PigGPA source code, analytical skills, benchmark scripts, scoring criteria, controlled verification tests and representative execution archives are available at https://github.com/ChongxiaoLiang/PigGPA. The PigGPA web service is hosted at https://pigga.zishuailab.com/.

